# Synonymous polymorphism difference relating to codon degeneracy between co-transcribed genes in the genome of *Escherichia coli*

**DOI:** 10.1101/2022.07.25.501341

**Authors:** Pratyush Kumar Beura, Piyali Sen, Ruksana Aziz, Chayanika Chetia, Madhusmita Dash, Siddhartha Shankar Satapathy, Suvendra Kumar Ray

## Abstract

The previous findings suggest that replication and transcription are two major reasons behind the different substitution patterns of mutations in genomic DNA. In the current work, we have compared the adjacent co-transcribed gene pairs regarding synonymous polymorphism in five different operons in *Escherichia coli*. It is interesting that the co-transcribed genes were different from each other regarding the polymorphism spectra. The transition to transversion ratio between gene pairs were different due to their compositional differences regarding two-fold degenerate codon and four-fold degenerate codons. Further, the polymorphism spectra difference between the gene pairs was more prominent in four-fold and six-fold degenerate codons than in the two-fold degenerate codons. In case of *rpoB* and *rpoC*, the major difference was found at UCC, GUA, CCG, GCU, GGC and CGC codons. Similarly, in case of the other four pairs of co-transcribed genes, the difference was more prominent in the higher degenerate codons than the two-fold degenerate codons. It may be that the restriction of two-fold degenerate codons to transition substitutions only regarding synonymous polymorphism is making these codons different from the higher degeneracy codons in this study.

## Introduction

Base substitution mutation is a major event of molecular evolution in organisms, which is known to be influenced by different factors such as DNA replication mechanism, damages in DNA such as cytosine deamination and guanine oxidation^1,2^ expression level of genes, recombination etc. It is well known that the asymmetry in DNA replication results different mutation patterns between the leading and the lagging strands in genomes resulting the former enriched with the keto nucleotides and the latter enriched with the amino nucleotides in bacteria^3,4^. In addition, genes near the origin of replication exhibits different mutation patterns than that at the terminus of replication: in bacteria the replication terminus region in a chromosome is known to be AT enriched in comparison to the origin of replication^5^. The role of transcription in causing mutation asymmetry in genes resulting higher C→T changes in the non-template strand than that in the template strand has been described recently^6^. Therefore, expression level of genes has different impact on mutation rates^7-9^. In addition to replication, localization, transcription, two genes can be different from each other regarding the amino acid level selection on their protein structure in a genome. Because of these complexities, it becomes difficult to compare between two genes within a genome regarding their polymorphism patterns. It is important to accurately compare mutation pattern between two genes because it helps to understand further the role intrinsic factors in polymorphism difference between the genes, if any.

In bacteria, one advantage to study molecular evolution is that two genes co-transcribed in an operon are adjacent to each other that are often located in the same strand and are similar regarding their gene expression at transcription level. The genes are often related in their function. Therefore, mutation pattern of two adjacent genes in an operon are likely to be similar, unless there are some unknown factors are causing mutation and/or selection biases in these genes.

In this study, our endeavour through this communication is to do the comparative analysis of mutation spectra in two adjacently placed co-transcribed genes in *Escherichia coli* genome. We have considered five pairs of cotranscribed genes that are present at different loci in *E*.*coli* genome. We have considered 5 pairs of genes (*rpoB/C, lacZ/Y, kdpA/B, araB/A* and *bcsA/B*) for the study (figure 1). It is interesting that two genes in any of these operons are found to be significantly different from each other regarding their synonymous polymorphism pattern. This suggests that variation arising in the genes are not identical to each other in spite of being similar regarding replication, transcription and strand localization. Understanding the mechanisms of this difference will expand our knowledge regarding the factors contributing towards the polymorphism spectra in genes. So far we understand this is the first such report of comparison of polymorphism spectra between two cotranscribed genes in bacteria.

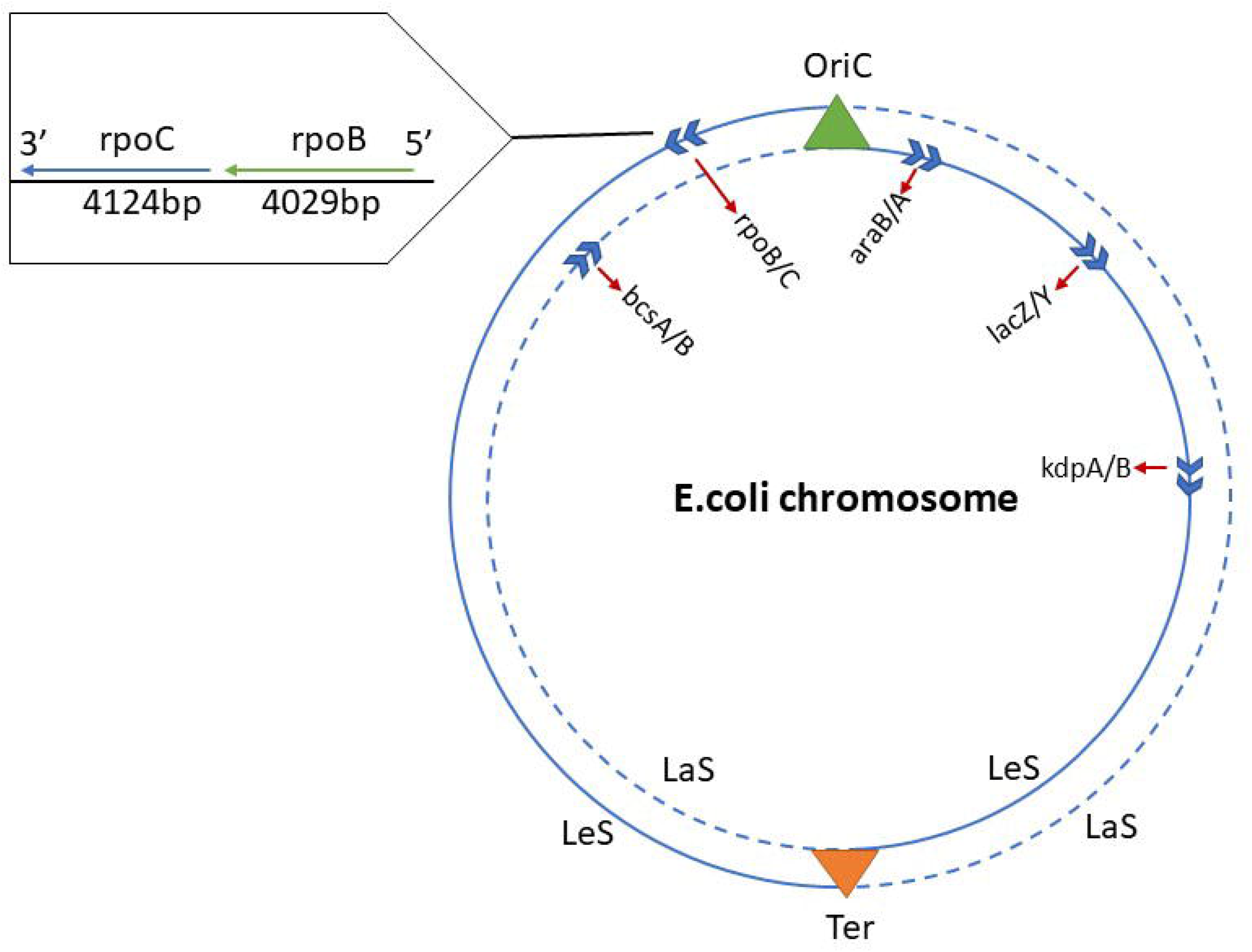

## Materials and Methods

### Selection of co-transcribed gene pairs from available dataset

The current study has considered co-transcribed genes in the *E. coli* (*E*.*coli*) genome for comparative mutational analysis between adjacent genes. Each pair of genes are co-transcribed in an operon and have related functions in cellular metabolic pathways. We started the work in 157 strains of *E. coli* bacterium for which alignments are available in public database^10^. The criteria of the selection of genes were set (>1200bp or >400 codons) in an assumption of getting considerable numbers of synonymous substitutions in each gene. We have selected 5 pairs of co-transcribed genes with different expression levels based on the above criteria. Out of 5 pairs of co-transcribed genes, 4 were in the leading strand (*LeS*) and only one (bcsA/B) was present in the lagging strand (*LaS*). The list of genes, as well as detailed related information, are enclosed in Table 1.

**Table 1.** Operons selected for the work, their coordinate and function details^1^.

| Operon | Genes | Start | End | Strand | Function |
| --- | --- | --- | --- | --- | --- |
| rpoBC | rpoB | 4181245 | 4185273 | LeS | DNA-directed RNA polymerase subunit beta |
|  | rpoC | 4185350 | 4189573 | LeS | DNA-directed RNA polymerase subunit beta' |
| lacZY | lacZ | 363231 | 366305 | LeS | beta-D-galactosidase |
|  | lacY | 361926 | 363179 | LeS | galactoside permease |
| kdpAB | kdpA | 727059 | 728732 | LeS | potassium-transporting ATPase subunit A |
|  | kdpB | 724988 | 727036 | LeS | potassium-transporting ATPase subunit B |
| araBA | araB | 68348 | 70048 | LeS | ribulokinase |
|  | araA | 66835 | 68337 | LeS | L-arabinose isomerase |
| bcsAB | bcsA | 3692618 | 3695236 | LaS | cellulose synthase catalytic subunit |
|  | bcsB | 3690268 | 3692607 | LaS | cellulose synthase periplasmic subunit |

### Derivation of reference sequence and finding the overall synonymous spectra

To find out the mutation pattern we first derived the reference sequence by comparing a gene sequence across the 157 strains. This procedure was followed as per the methodology developed in our lab^11^. Strains having ‘N’ were not considered for the study. We maintain a common set of strains for each pair of genes in the study. The reference sequence was derived for each gene by considering the most frequent nucleotide present in a certain position in the alignment as a reference nucleotide for the position. The compositional details of the reference sequences for all the genes were calculated. AT skew was calculated by (A-T)/(A+T) and GC skew was calculated by (G-C)/(G+C). RY and KM skew was also calculated to know the inequality of purine: pyrimidine and Keto: Amino inequality in genes. RY skew was calculated as [{(A+G) - (T+C)} / (A+T+C+G)] and KM skew was calculated as [{(T+G) -(A+C)}/ (A+T+C+G)] (Table-2). Methods of finding mutations is explained in Supplementary Table 1. Only synonymous mutations were considered in this study. The number of mutations were normalized by dividing the number of mutations by the total number of nucleotides present in the gene. Suppose we found 30 C→T substitutions and the number of C nucleotides in the gene is 150. Then the normalized C→T mutation frequency is calculated as 30/150=0.2. We estimated normalized mutation spectra of all the twelve substitutions for the five pairs of genes and presented in result section.

**Table 2.** composition and skew detailed information of gene pairs.

| Gene | A | T | C | G | SIZE | GC% | AT | GC | RY | KM |
| --- | --- | --- | --- | --- | --- | --- | --- | --- | --- | --- |

<sup>1</sup> Names of the 5 pairs of genes as well as the strand, start and end coordinates and function based on data availability in National Center of Biotechnology Information (NCBI) by considering the reference strain *Escherichia coli* str. K12 substr. MG1655 (NCBI: txid511145)

|  |  |  |  |  |  |  | SKEW | SKEW | SKEW | SKEW |
| --- | --- | --- | --- | --- | --- | --- | --- | --- | --- | --- |
| *rpoB* | 1016 | 891 | 1034 | 1088 | 4029 | 52.67 | 0.066 | 0.025 | 0.044 | -0.018 |
| *rpoC* | 1010 | 938 | 1090 | 1186 | 4224 | 53.88 | 0.037 | 0.042 | 0.040 | 0.006 |
| *lacZ* | 677 | 665 | 845 | 888 | 3075 | 56.36 | 0.009 | 0.025 | 0.018 | 0.010 |
| *lacY* | 239 | 433 | 284 | 298 | 1254 | 46.41 | -0.289 | 0.024 | -0.144 | 0.166 |
| *kdpA* | 293 | 451 | 425 | 505 | 1674 | 55.56 | -0.212 | 0.086 | -0.047 | 0.142 |
| *kdpB* | 433 | 465 | 533 | 618 | 2049 | 56.17 | -0.036 | 0.074 | 0.026 | 0.057 |
| *araB* | 349 | 361 | 489 | 502 | 1701 | 58.26 | -0.017 | 0.013 | 0.001 | 0.015 |
| *araA* | 340 | 328 | 416 | 419 | 1503 | 55.56 | 0.018 | 0.004 | 0.010 | -0.006 |
| *bcsA* | 526 | 638 | 704 | 751 | 2619 | 55.56 | -0.096 | 0.032 | -0.025 | 0.061 |
| *bcsB* | 518 | 509 | 618 | 659 | 2304 | 55.43 | 0.009 | 0.032 | 0.022 | 0.014 |

### Finding the ratio of transition to transversion in gene pairs

Transition (Ti) to transversion (Tv) ratio of all the genes were calculated. Overall, Ti/Tv of the gene and Ti/Tv at four-fold degenerate site were calculated for each gene to show the variation between cotranscribed genes. For overall synonymous transition/synonymous transversion (STi/STv), the values were collected from the initial non-normalized synonymous spectra table (Supplementary table 2). For four-fold degenerate site Ti/Tv, the Ti and Tv values were calculated by considering as synonymous substitutions at four-fold sites for individual genes. for each gene the summation of 5 four-fold degenerate sites mutation was taken to get the Ti/Tv values at four-fold degenerate sites. This method was followed for other genes. Further, four-fold degenerate individual amino acids were also considered individually to find out the Ti/Tv ratio per amino acids and then comparison between two co-transcribed genes were done under this criterion.

### Mutational frequency comparison at the codon level

From the reference sequence of each gene, we computed for codon count using the web portal http://agnigarh.tezu.ernet.in/~ssankar/cbb_tu.html12. Generally, the analyses and comparisons were done for each pair of genes individually. We started the work from *rpoB* and *rpoC* gene, after getting the codon count of both separate analyses for four-fold codons (20 codons), two-fold codons (18 codons), and six-fold codons (split box and family box) were performed. Four-fold codons are such codons in genetic codon table where single amino acid is coded by 4 codons in a family box, like Valine is coded by GUN. Similarly, two-fold are coded by only 2 codons in the codon table like GAY/GAR etc. The synonymous mutations involved in the 3^rd^ position of codons were calculated for each box separately. Suppose we got three synonymous mutations in 3^rd^ position of UUU in *rpoB*, a value of 3 was entered in the mutation column against UUU for *rpoB*. Similarly, synonymous mutations in UUU of *rpoC* was also mentioned in the adjacent box for comparison Then, mutation frequency was calculated by taking the number of synonymous mutations involved in 3^rd^ position of a certain codon and dividing it with the number of that codon present in the gene. This technique was followed for the remaining pair of genes. In Supplementary table 3 a detailed procedure of codon count, mutation and frequency of *rpoB* and *rpoC* is provided. The criteria of consideration of mutations were set as codon abundance >30 or mutation frequency >5 per codon. The mutation frequency showing the difference on the basis of the said parameter was noted. Finally, four-fold degenerate (FFD) codons, two-fold degenerate codons (TFD), six-fold degenerate codons (SFD) mutation frequency information of a pair of gene was compared through a Box whisker plot, generated through Python script. SFD amino acid (L, S, R) codons are divided into 2 segments: family box (CUN, UCN, CGN) and split box (UUR, AGY, AGR). SFD family box was only considered to calculate synonymous mutation in 3^rd^ position due to presence of less numbers of mutations. This step was performed to show the box-wise mutation difference between adjacent genes. All the statistical methods used in the study are mentioned in the result section of the text.

## Results

### The synonymous polymorphism spectra difference between co-transcribed genes

Two adjacent co-transcribed genes in a bacterial chromosome are similar in various aspects such as position in chromosomes, strand localization, and transcriptional expression level. We studied *rpoB/C* and *kdpA/B* gene pairs and quantified their polymorphisms difference in individual strain (Supplementary table 7). For example, in 4 strains of *E*.*coli* we found, 0 substitutions in *rpoB* and 17 substitutions in *rpoC*. There were only 5 strains in *E*.*coli* where both the *rpoB* and *rpoC* were having similar numbers of substitutions. In a strain in *E*.*coli* we found, 52 substitutions in *kdpA* and 22 substitutions in *kdpB*, similarly in one more strain it was 52 substitutions in *kdpA* and 48 substitutions in *kdpB*. In the plot of substitution difference between *rpoB* and *rpoC* of individual strains (*rpoB-rpoC*), we found minimum substitution difference as −16 to 19. Whereas, in case of *kdpA* and *kdpB* (*kdpA-kdpB*) it was found to be −23 to 30. It was evident that the co-transcribed genes were different from each other even at the individual strain level. Therefore, we compared the genes regarding their synonymous polymorphisms (Table 3). We have plotted a graph showing the individual strain wise mutations in both the gene pairs. The graph clearly shows the differences between adjacently placed co-transcribed genes even at individual strain level (Supplementary table 7). In case of *rpoB* and *rpoC*, the two genes were different from each other regarding C→T and G→A polymorphisms: in *rpoB* and *rpoC*, C→T frequency was 0.074 and 0.058 respectively, while G→A frequency was 0.016 and 0.024, respectively. It is pertinent to note that C→T frequency was about ∼five-fold higher than G→A in *rpoB* whereas in *rpoC*, C→T frequency was only about two-fold higher than G→A. This observation indicated that the two co-transcribing genes were not similar regarding synonymous transition polymorphisms. In another pairs of genes such as *kdpA* and *kdpB*, the A→G frequency was 0.051 and 0.028, respectively; in *araB* and *araA*, the T→C frequency was 0.053 and 0.098 respectively; in *bcsA* and *bcsB*, the G→A frequency was 0.060 and 0.049, respectively. C→T frequency was the highest among polymorphism spectra in all the genes. It is interesting that all the five pairs under the study exhibited difference from each other. We could not observe any pattern conserved across different gene pairs regarding their polymorphism differences such as *rpoB* and *rpoC* were prominently different regarding C→T, whereas *lacZ* and *lacY* were prominently different regarding T→C. In future it will be an interesting to find out the reason for these differences among the different gene pairs.

**Table 3.**
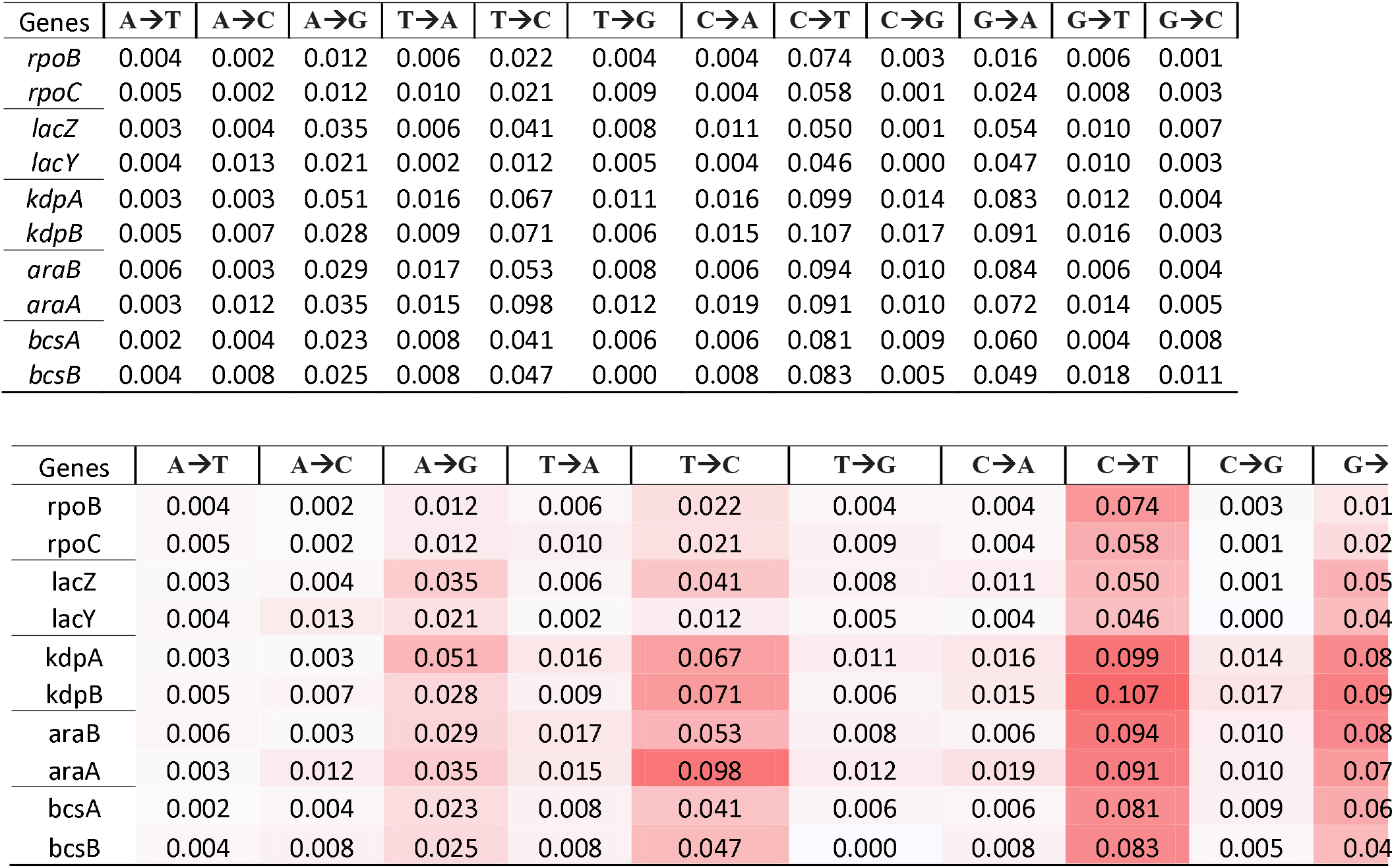
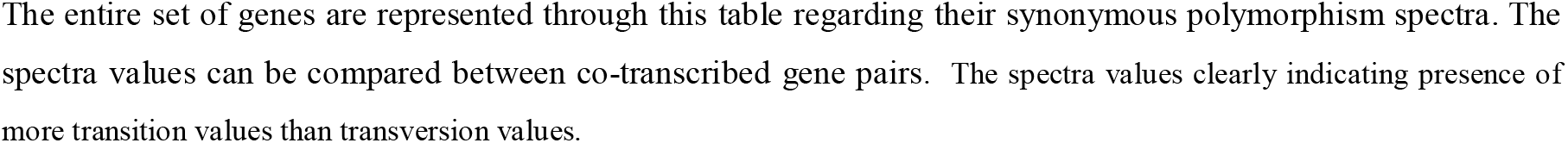
Synonymous mutational spectra of genes are presented in tabular form, the synonymous spectra values in pair of co-transcribed genes can be compared here.

### Ti/Tv ratio difference between the co-transcribed genes due to codon degeneracy composition difference

We studied synonymous transition to synonymous transversion ratio in all genes (Table 4). It was evident that the two cotranscribed genes were different from each other regarding the ratio as follows: in *rpoB* the value was 4.33 whereas in *rpoC* it was 2.9; in *araB* the value was 4.9 whereas in *araA* it was 3.1, in *bcsA* the value was 4.3 and in *bcsB* the value was 3.3. This indicated that Ti/Tv ratio are different among the genes. We studied Ti/Tv at FFD in all these 10 genes. Ti/Tv values at FFD were observed to significantly lower (*p* value < 0.01; Mann Whitney U test) than that in the whole gene because of difference between Ti and Tv rate in organism (Supplementary Table 4). It is pertinent to note that synonymous polymorphisms at TFD sites are only possible due to transitions whereas the same at FFD sites are possible by both transition and transversion. We found out the TFD:FFD ratio in these 10 genes. We did a correlation between the TFD:FFD ratio with the difference between Ti/Tv values between whole gene and at FFD. The strong positive correlation (*Pearson r* value 0.668) suggested that higher the composition of TFD in a gene greater will be Ti/Tv ratio difference between the whole gene and the FFD. Therefore, the compositional difference between TFD and FFD among the genes influences synonymous Ti/Tv ratio in genes. We studied the Ti/Tv ratio at four-fold degenerate sites at individual amino acid level (Supplementary table 5). The two co-transcribed were found to be distinctly different from each other in certain amino acid codons: in case of Gly, Ti/Tv ratio in *rpoB* and *rpoC* was 1.75 and 6.50 respectively; in case of Ala, Ti/Tv ratio in *lacZ* and *lacY* was 1.27 and 4.00 respectively.

**Table 4.**
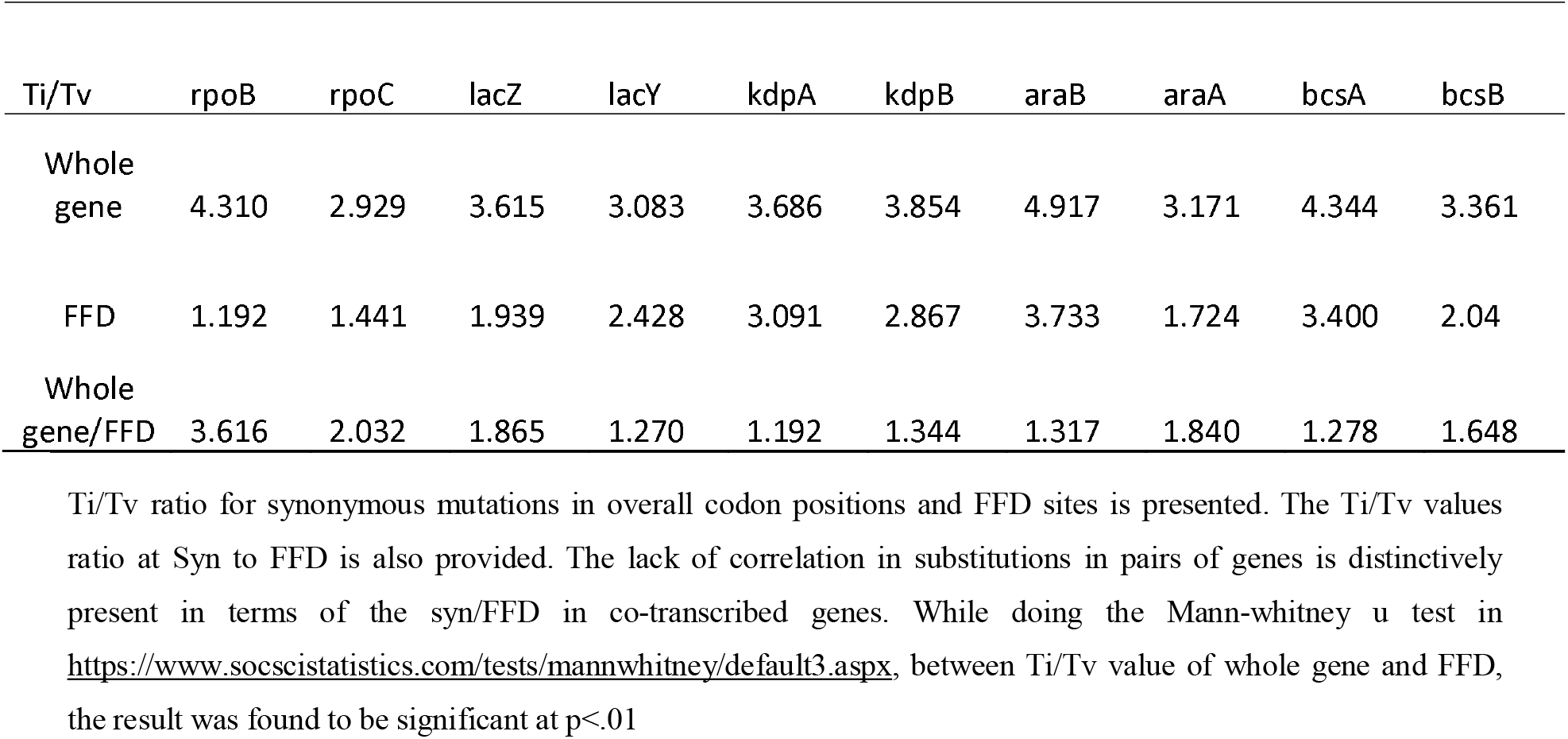
Ti/Tv ratio comparison of 5 pairs of genes in terms of overall synonymous and FFD site.

### Synonymous polymorphism difference between co-transcribed genes at four-fold degenerate sites, two-fold degenerate sites and six-fold degenerate (family box) codons

Considering the rate of a transition four times more frequent than that of a transversion, the synonymous sites value in a FFD codon is 1.000 whereas, synonymous site value in a TFD codon is 0.667^13^. Therefore, synonymous polymorphisms at a FFD site are expected to be 1.5 times more frequent than that at a TFD site. To compare synonymous polymorphism at TFD and FFD, we studied synonymous polymorphism in each codon. In supplementary Table 6 synonymous polymorphisms involved at the 3^rd^ position of codons is presented. Codons such as GUA, CCG, GCU, GGC synonymous polymorphisms were different between *rpoB* and *rpoC*. The codon occurrence of GUA was 31 and 32 for *rpoB* and *rpoC*, the synonymous polymorphisms recorded as 3 and 7 for *rpoB* and *rpoC* respectively, hence the frequency noted as 0.097 and 0.219 respectively for *rpoB* and *rpoC*. Here it can be vindicated that, a codon having similar abundance in both adjacent genes had more than 2-fold polymorphism frequency difference. The codon occurrence of GCU was noted as 19 for *rpoB* and 28 for *rpoC*, the synonymous substitutions were noted as 1 for *rpoB* and 7 for *rpoC*, hence frequency was found to be 0.053 for *rpoB* and 0.250 for *rpoC*, which was ∼5-fold frequency difference between both the genes in GCU codon. Similarly, the codon occurrence of CCG was found to be 38 for *rpoB* and 45 for *rpoC*, with substitutions recorded as 3 and 8 for *rpoB* and *rpoC*, respectively and frequencies were 0.079 and 0.178. The codon occurrence of GGC was 35 in *rpoB* and 29 in *rpoC*, with synonymous substitutions as 4 in *rpoB* and 9 in *rpoC*, with a frequency of 0.114 in *rpoB* and 0.310 in *rpoC*.

The differences in patterns observed in primary synonymous spectra was not present universally in our work. For example, we have found differences in *rpoB*/*C* gene C→T spectra in UUC and CGC (six-fold; family box) codons, whereas C ending codons in TFD and FFD codons show no difference. Whereas the difference in T→C spectra observed in initial synonymous spectra of *araB*/*A* was not found in six-fold U ending codons (family box), rather CAU and GGU codons show the difference.

Among six-fold degenerate (family box) codons the prominent difference was noted in CUG, UCC and CGC codons. The codon occurrence in CUG was found to be 100 for *rpoB* and 125 for *rpoC*, with synonymous substitutions as 6 in *rpoB* and 13 in *rpoC*, with frequency as 0.06 and 0.104 respectively. Similarly, the codon occurrence of UCC was found to be 31 in *rpoB* and 27 in *rpoC*, with synonymous substitutions as 9 in *rpoB* and 2 in *rpoC*, with a frequency of 0.290 in *rpoB* and 0.074 in *rpoC*. Codon occurrence of CGC was found to be 28 in *rpoB* and 24 in *rpoC*, with synonymous substitutions as 10 in *rpoB* and 1 in *rpoC*, with a frequency of 0.357 in *rpoB* and 0.042 in *rpoC*. Regarding TFD, only one codon, CAC was exhibiting noticeable difference: it had an occurrence of 18 in *rpoB* and 17 in *rpoC*, but synonymous substitution as 5 in *rpoB* and 0 in *rpoC*, 0.278 and 0 mutation frequency for *rpoB* and rpoC respectively.

Out of the 12 codons belonging to the three family boxes of Leu, Ser and Arg, only three codons were exhibiting the difference. Out of the 20 codons belonging to the five family boxes in case of Val, Ala, Pro, Thr, Gly, only four codons were exhibiting the difference. In Table 5 mutation frequency comparison of 5 pairs of co-transcribed genes and four-fold, two-fold and family box of six-fold amino acids are presented. Similarly, *lacZ*/*Y* was showing significant frequency difference at 4 four-fold codons; CCG, ACA, GCG and GGU. *kdpA*/*B* were showing frequency differences at 4 codons: GUG, GCC, GGC and GGG.

**Table 5.**
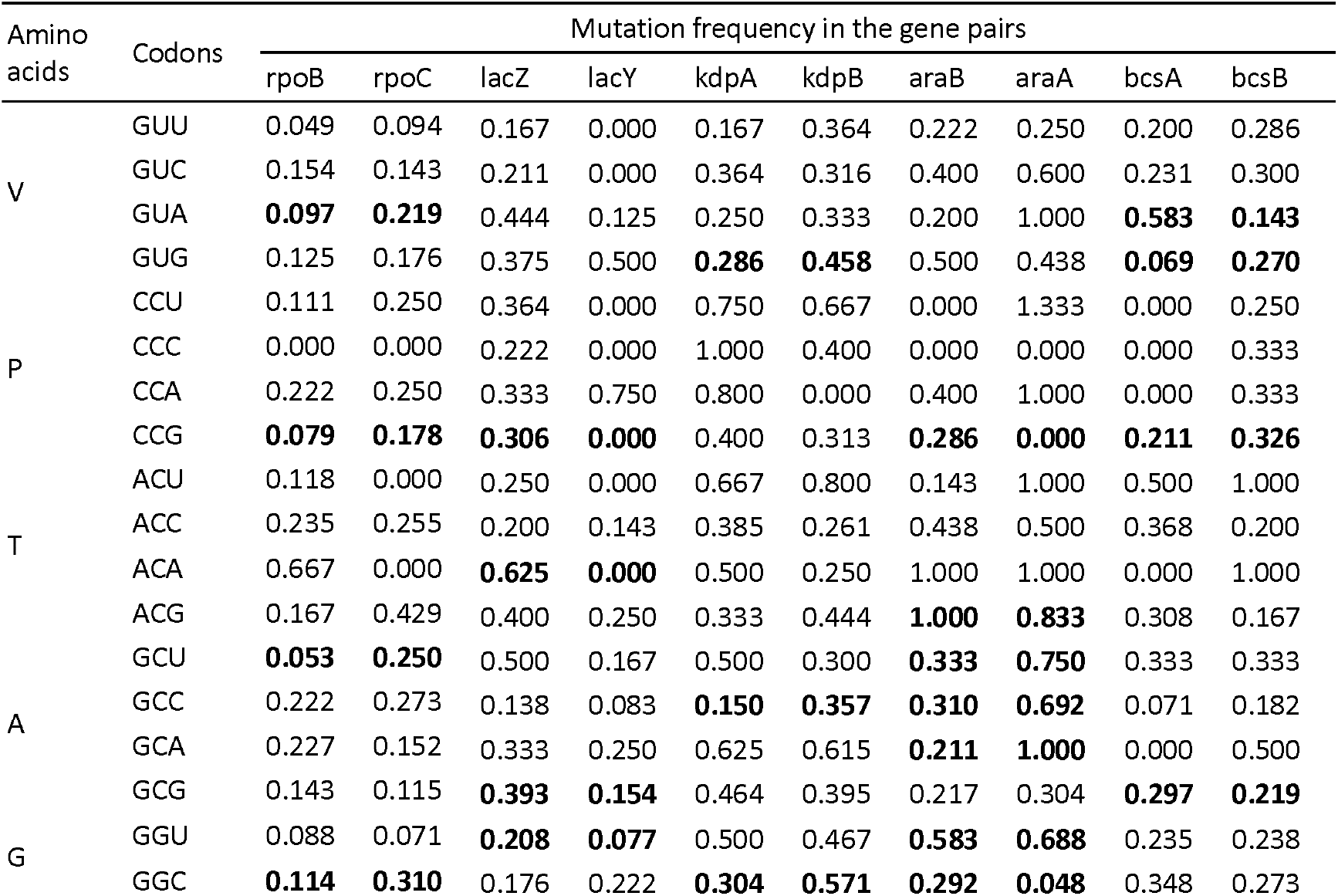
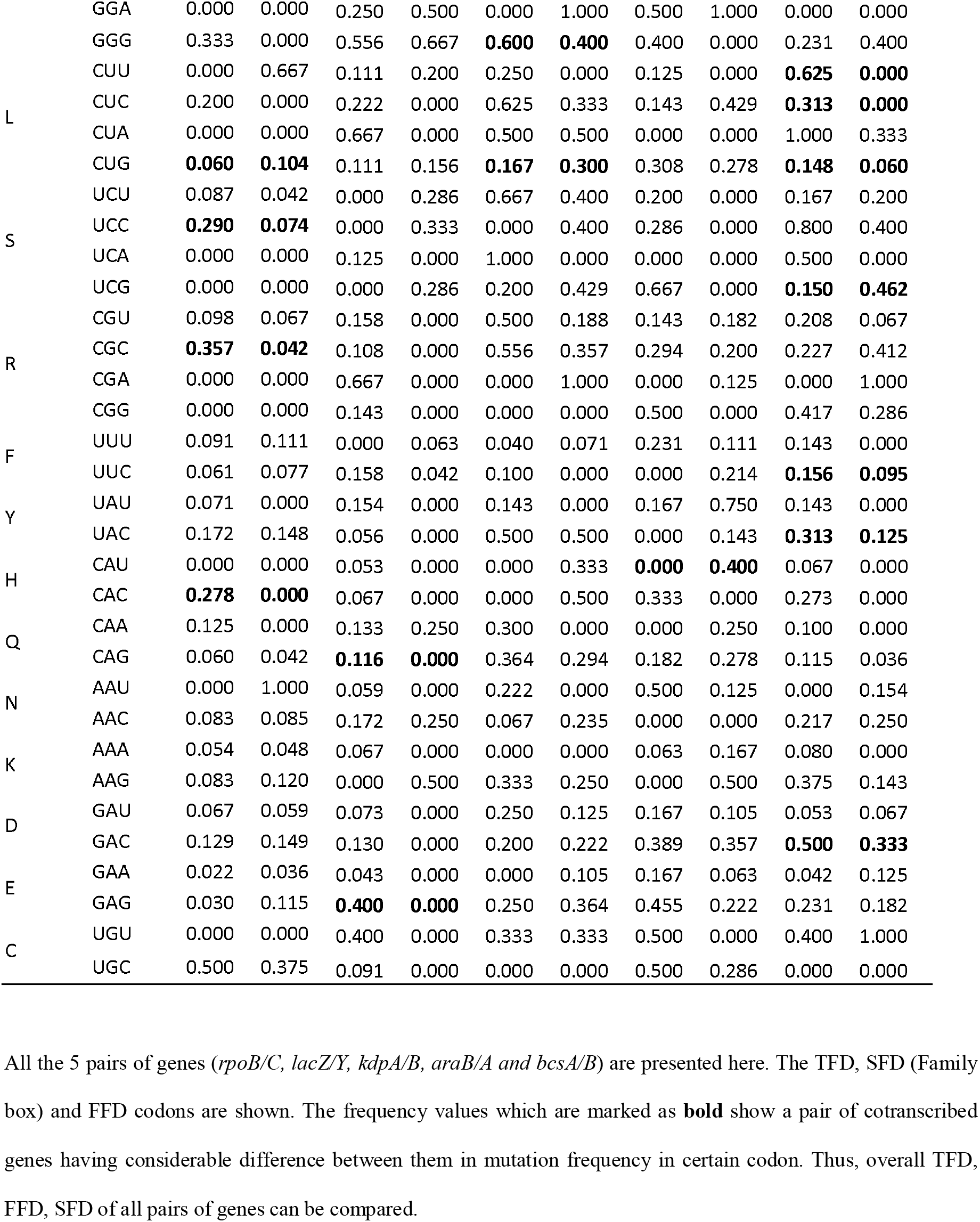
mutation frequency comparison at codon specific level of all genes.

Likewise, we found out difference between *araB*/A at 7 nos. of four-fold codons, and between *bcsA/B* pair at 4 nos. of four-fold codons. Among 5 pairs, *bcsA/B* were showing difference at 3 codons in two-fold codons. whereas other pairs were not exhibiting such differences at two-fold codons. Till now, it was evident that four-fold codons were predominantly showing differences in all gene pairs. For entire 5 pairs of genes, the quartile ranges as well as the median values in the box-plot for four-fold codons were showing differences. Whereas in two-fold codons only *lacZ/Y* and *bcsA*/*B* were showing differences in both quartile range and median values. As six-fold codons were representing only family box substitutions, gene pairs were showing the difference in box-plot regarding the median and quartile range. The box-plots were clearly indicating the fold wise differences between co-transcribed gene pairs, out of which differences at four-fold codons were principally found between a pair of genes. Further we have done a Mann-Whitney u test between entire FFD and TFD, and between entire FFD and SFD and the result were found to be significantly different at *p*<0.01. In an interesting comparison between GAU and GGU codons of *araB/A* we got to know, the codon occurrence for GAU in *araB*/*A* was found to be 12 and 19 respectively, whereas 2 mutations were found per each gene indicating a mutation frequency of 0.167 and 0.105 respectively. While, in GGU, the codon occurrence was found to be 12 and 16 respectively, whereas 7 and 11 numbers of mutations were recorded, indicating a frequency of 0.583 and 0.688 respectively. This was a big evidence of high mutation frequency of four-fold codons which was observed in many cases though.

## Discussion

In the present study five different sets of co-transcribed genes have been compared regarding synonymous polymorphism in *E. coli*. The adjacent genes are observed to be different regarding synonymous polymorphism. Though the two adjacent genes in an operon are similar regarding replication, strand location and transcript level expression, their difference regarding synonymous polymorphism is interesting. TFD codons can undergo synonymous polymorphism only by transition substitution while the higher degenerate codons can undergo synonymous polymorphism by both transition as well as transversion substitutions. The rate of a transition substitution is four times more than that of a transversion substitution^14^. and the purifying selection is stronger on non-synonymous substitutions than synonymous substitutions. These cause higher transversion proportion in FFD than TFD codons. Therefore, Ti/Tv is higher is genes having lower proportion of FFD codons across the five gene pairs in this study. The current study has manifested the role of codon degeneracy due to the difference in mutation spectra of two co-transcribed genes. Yet, among all, FFD codons remained to be the common cause behind the mutational spectra difference in each gene pair. In case of the higher degeneracy codons synonymous transversion is possible unlike the two-fold degenerate codons. Whether this is the only reason for the difference or there are additional reasons involves will be interesting to find out in future studies. The observation regarding codon degeneracy and difference in polymorphism between adjacent genes can be analysed in the light of codon context. It is highly improbable that context of TFD codons remain invariable between the genes while the context of higher degenerate codons is variable between the genes. Considering that codon context is variable between the adjacent genes both in case of TFD codons as well as FFD codons, codon context might not be a potential reason for the difference between the two genes.

It has already been reported by the researchers that synonymous polymorphisms though do not affect the amino acid sequence in a protein, it influences its function by protein folding. Whether TFD codons and FFD codons attribute differently to protein folding is yet to be discovered. The difference between the degenerate codons invokes many fundamental questions in genetic code evolution. Degeneracy to amino acid codons have been assigned randomly or degeneracy has a role in protein folding according to which degeneracy have been assigned to amino acids. Future research will shed more light on it.

## Supporting information

Figure 2. Box plot showing mutation frequency of gene pairs in an FFD/TFD/SFD manner

## Acknowledgments

PKB is grateful to Tezpur University for the institute fellowship. PS is grateful to UGC, GoI New Delhi, for the JRF. MD is grateful to NIT, Arunachal Pradesh. SSS and SKR are grateful to DBT, GoI, for the Computational biology and bioinformatics center at Tezpur University.

## Authors’ Contributions

Conceptualization: PKB, SSS and SKR; Methodology: PKB, PS, SSS and SKR; Writing, review and editing: PKB, PS, RA, CC, MD, SSS and SKR; Supervision: SSS and SKR. All authors endorsed the manuscript.

---

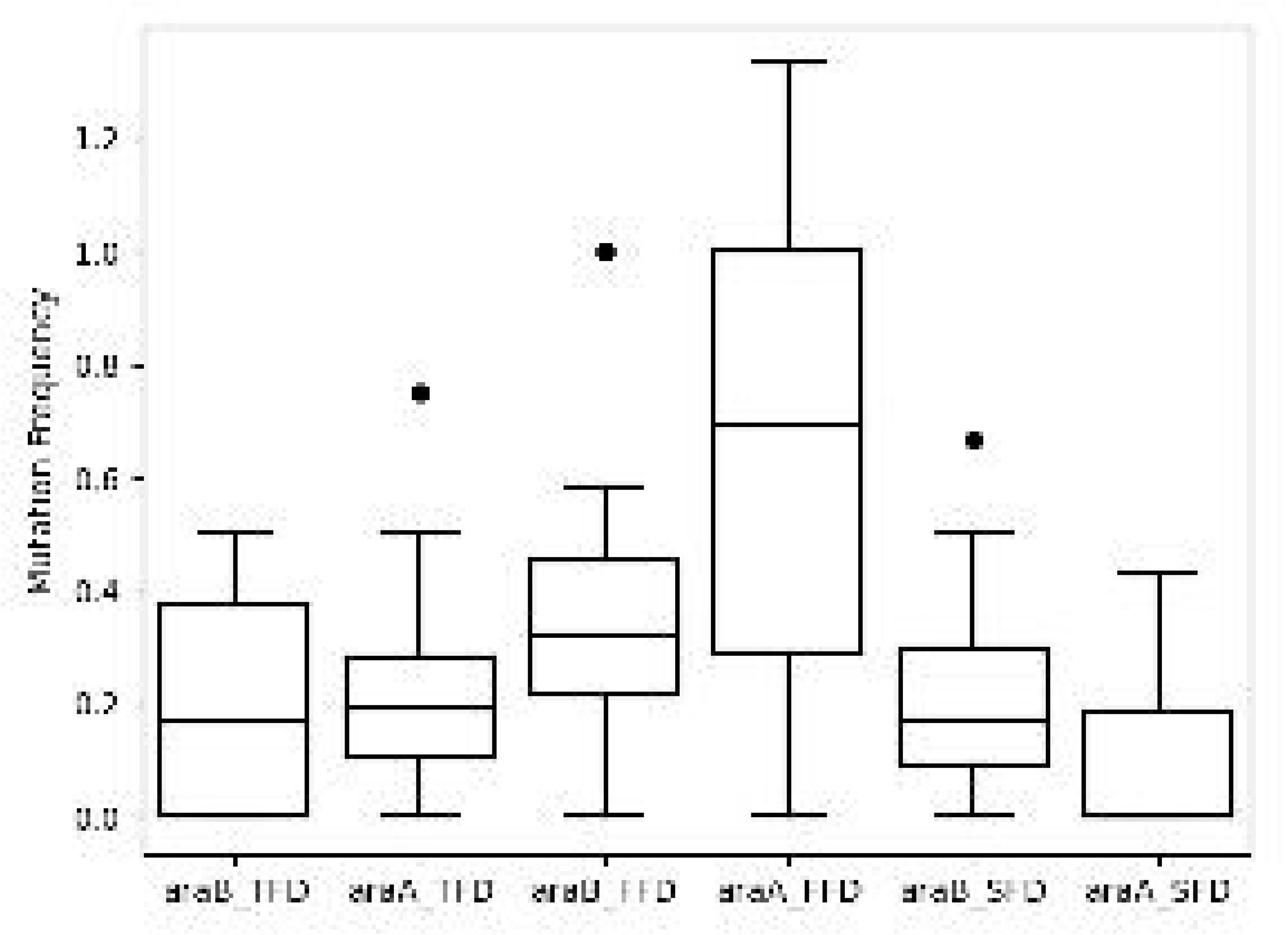

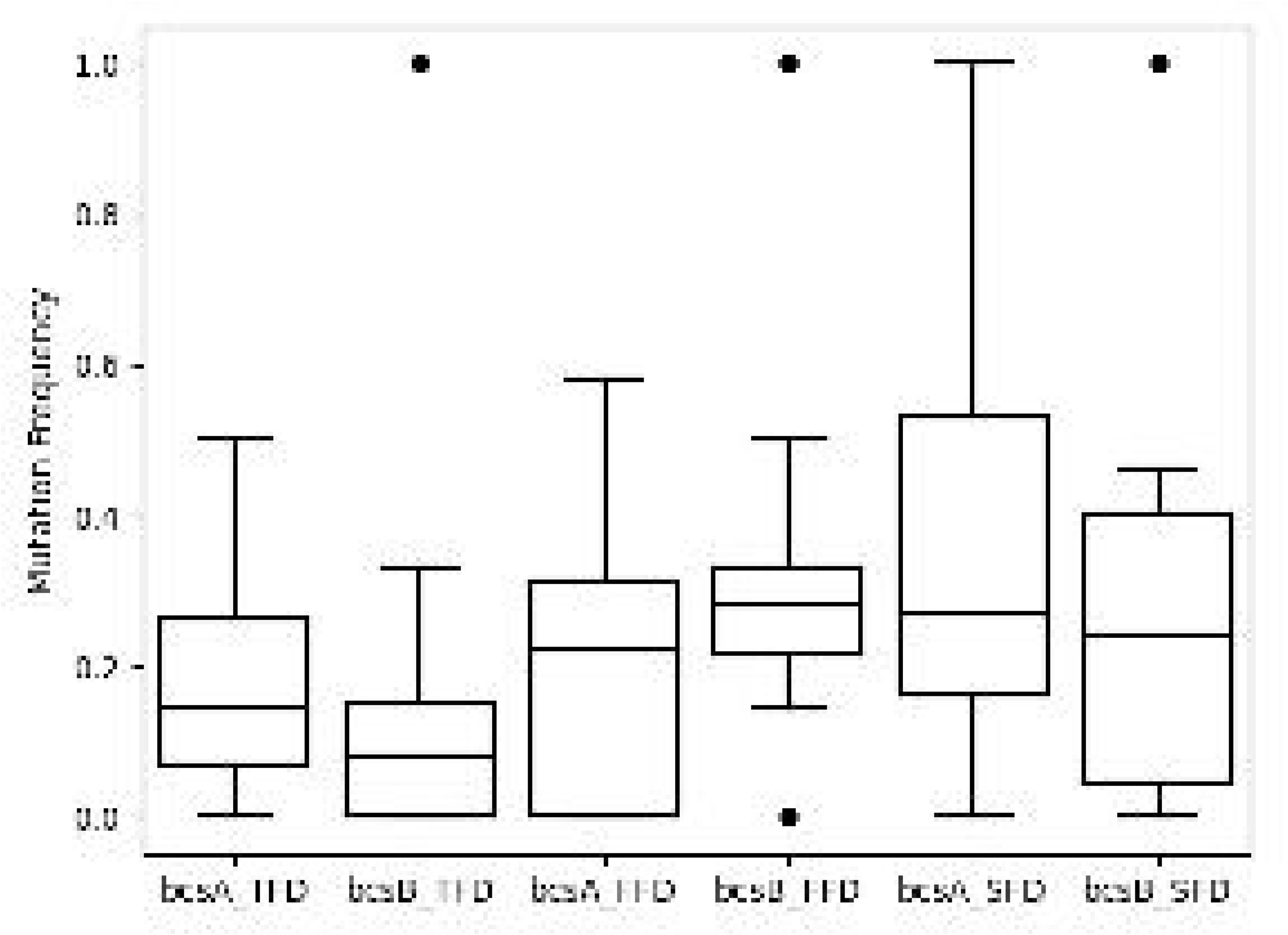

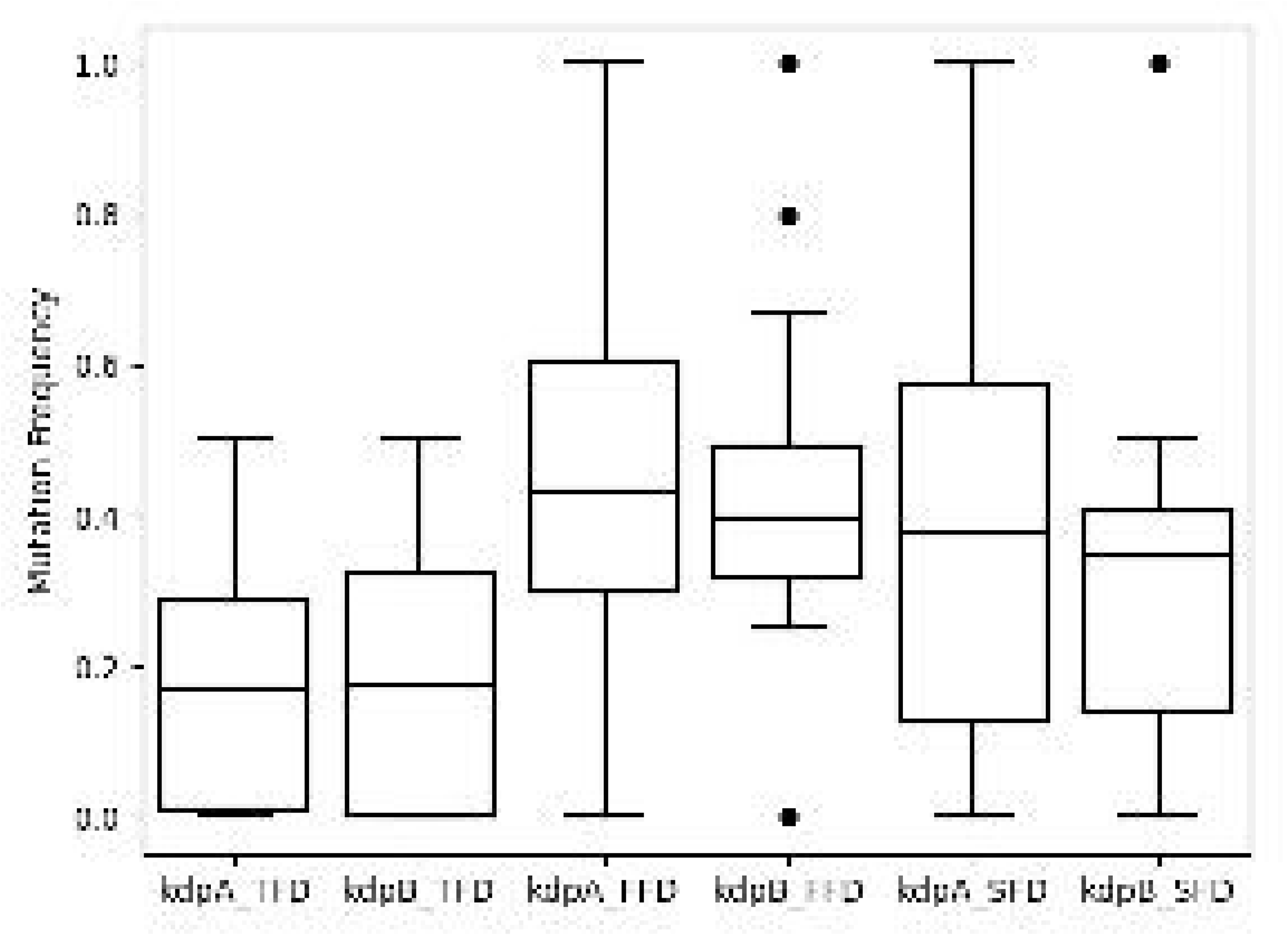

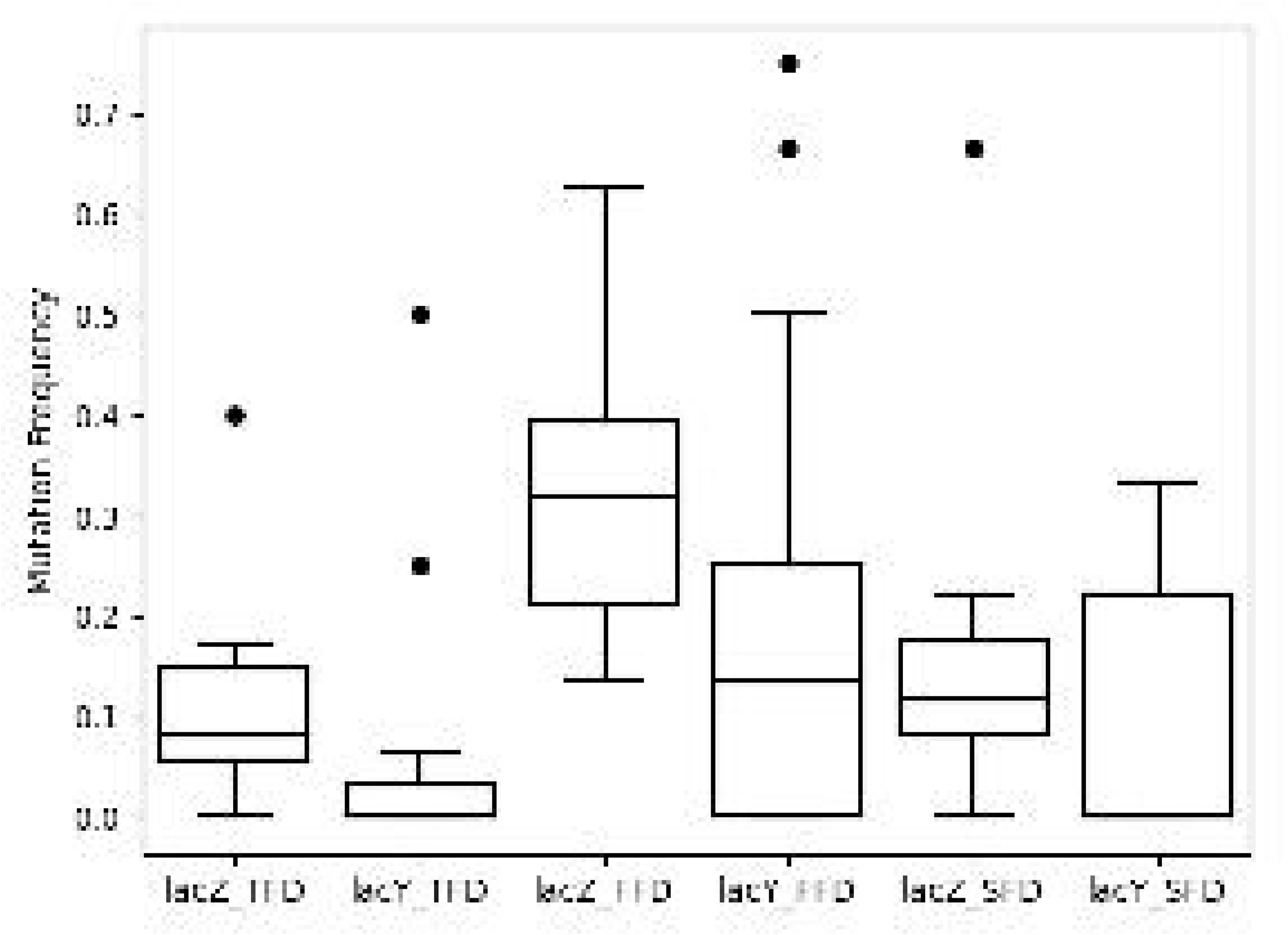

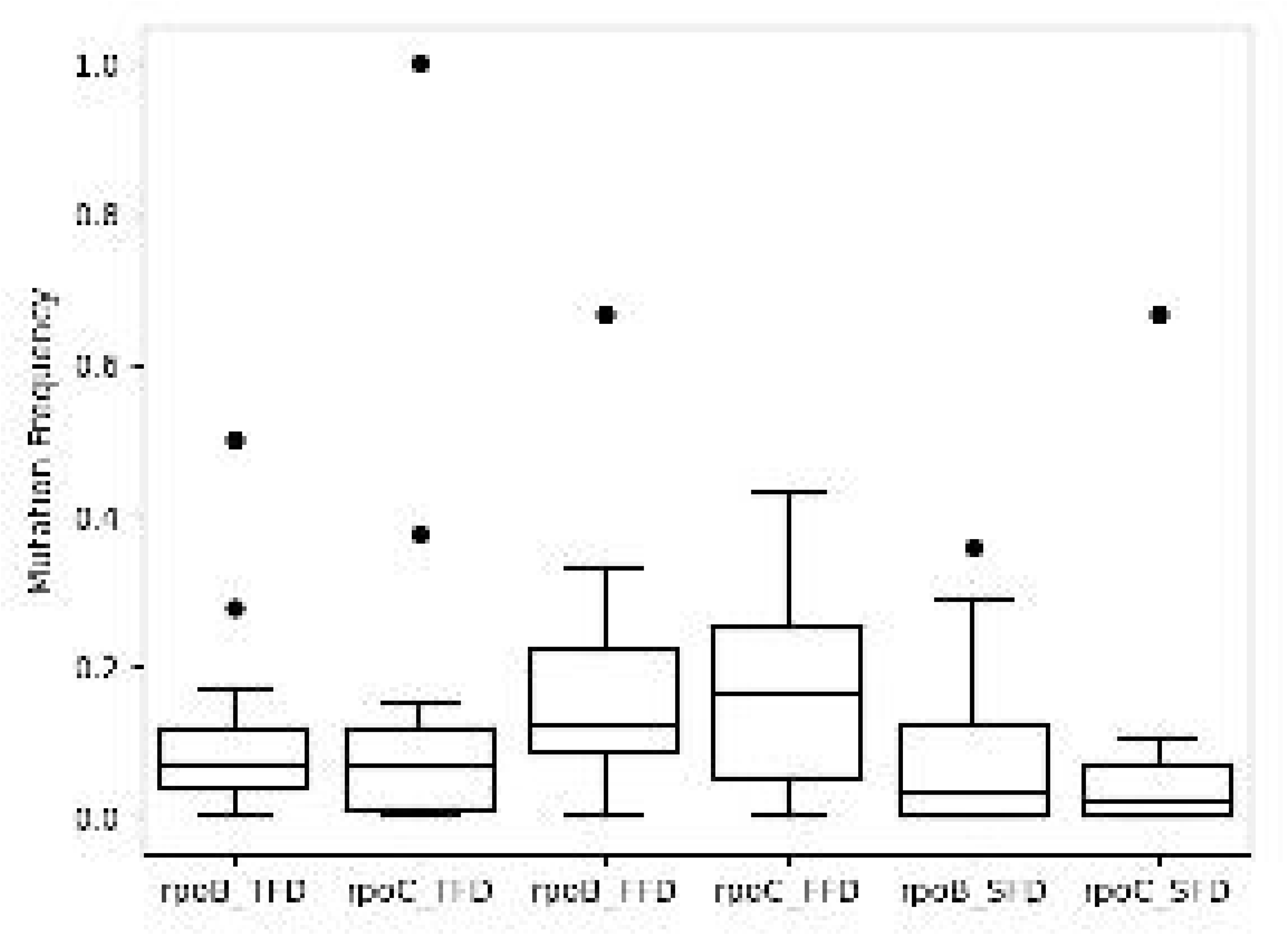

