## Supplementary material for "Synonymous polymorphism difference relating to codon degeneracy between co-transcribed genes in the genome of *Escherichia coli*": Figure 2. Box plot showing mutation frequency of gene pairs in an FFD/TFD/SFD manner

Current_science_supplementary

**Supplementary table 1. methodology for calculation of mutations.**

| **POSITION** | **1** | **2** | **3** | **4** | **5** | **6** | **7** | **8** | **9** |
| --- | --- | --- | --- | --- | --- | --- | --- | --- | --- |
| **ST1** | **A** | **A** | **T** | **A** | **G** | **C** | **C** | **T** | **A** |
| **ST2** | **A** | **A** | **T** | **A** | **G** | **T** | **C** | **A** | **A** |
| **ST3** | **A** | **A** | **T** | **A** | **G** | **A** | **C** | **A** | **A** |
| **ST4** | **A** | **A** | **A** | **A** | **G** | **G** | **C** | **A** | **T** |
| **ST5** | **A** | **A** | **A** | **T** | **G** | **G** | **C** | **A** | **T** |
| **ST6** | **A** | **A** | **T** | **T** | **G** | **G** | **C** | **A** | **T** |
| **ST7** | **A** | **A** | **T** | **T** | **G** | **G** | **C** | **A** | **A** |
| **ST8** | **A** | **T** | **G** | **T** | **G** | **G** | **C** | **A** | **A** |
| **ST9** | **A** | **T** | **G** | **T** | **G** | **G** | **T** | **A** | **T** |
| **ST10** | **A** | **T** | **G** | **T** | **G** | **G** | **T** | **A** | **T** |
| **A** | **10** | **7** | **2** | **4** | **0** | **1** | **0** | **9** | **5** |
| **T** | **0** | **3** | **5** | **6** | **0** | **1** | **2** | **1** | **5** |
| **C** | **0** | **0** | **3** | **0** | **0** | **1** | **8** | **0** | **0** |
| **G** | **0** | **0** | **0** | **0** | **10** | **7** | **0** | **0** | **0** |
| **Reference Sequence** | **A** | **A** | **T** | **T** | **G** | **G** | **C** | **A** | **?** |
| **MUTATION** |  | **A-->T** | **T>C, T>A** | **T>A** |  | **G>A, G>T, G>C** | **C>T** | **A>T** | **N/A** |

Hypothetical mutation scenario to explain the methodology that has been followed in this work. In position 1 A is found 10 times hence in Reference Sequence its written as A. similarly in 2^nd^ position A was found 7 times and T was found 3 times. Hence it can be said that the mutation was observed in A🡪T pattern. In the last position we found 5 numbers of A & T each, here we can’t assume which one can be taken in reference sequence, so we had to keep in mind the during the work

**Supplementary table 2. Spectra showing number of synonymous substitutions observed for each gene, before normalization**.

| Genes | A->T | A->C | A->G | T->A | T->C | T->G | C->A | C->T | C->G | G->A | G->T | G->C | TOTAL |
| --- | --- | --- | --- | --- | --- | --- | --- | --- | --- | --- | --- | --- | --- |
| rpoB | 4 | 2 | 12 | 5 | 20 | 4 | 4 | 76 | 3 | 17 | 6 | 1 | 154 |
| rpoC | 5 | 2 | 12 | 9 | 20 | 8 | 4 | 63 | 1 | 29 | 9 | 3 | 165 |
| lacZ | 2 | 3 | 24 | 4 | 27 | 5 | 9 | 42 | 1 | 48 | 9 | 6 | 180 |
| lacY | 1 | 3 | 5 | 1 | 5 | 2 | 1 | 13 | 0 | 14 | 3 | 1 | 49 |
| kdpA | 1 | 1 | 15 | 7 | 30 | 5 | 7 | 42 | 6 | 42 | 6 | 2 | 164 |
| kdpB | 2 | 3 | 12 | 4 | 33 | 3 | 8 | 57 | 9 | 56 | 10 | 2 | 199 |
| araB | 2 | 1 | 10 | 6 | 19 | 3 | 3 | 46 | 5 | 42 | 3 | 2 | 142 |
| araA | 1 | 4 | 12 | 5 | 32 | 4 | 8 | 38 | 4 | 30 | 6 | 2 | 146 |
| bcsA | 1 | 2 | 12 | 5 | 26 | 4 | 4 | 57 | 6 | 45 | 3 | 6 | 171 |
| bcsB | 2 | 4 | 13 | 4 | 24 | 0 | 5 | 51 | 3 | 32 | 12 | 7 | 157 |

**Supplementary table 3. Codon count, mutations and mutation frequency of rpoB & rpoC. this method was followed for remaining gene pairs.**

| Amino Acids | Codons | Codon count | | Mutations | | Mutation frequency | |
| --- | --- | --- | --- | --- | --- | --- | --- |
|  |  | rpoB | rpoC | rpoB | rpoC | rpoB | rpoC |
| V | GUU | 41 | 53 | 2 | 5 | 0.049 | 0.094 |
|  | GUC | 13 | 7 | 2 | 1 | 0.154 | 0.143 |
|  | **GUA** | **31** | **32** | **3** | **7** | **0.097** | **0.219** |
|  | GUG | 24 | 17 | 3 | 3 | 0.125 | 0.176 |
| P | CCU | 9 | 4 | 1 | 1 | 0.111 | 0.250 |
|  | CCC | 0 | 0 | 0 | 0 | 0.000 | 0.000 |
|  | CCA | 9 | 8 | 2 | 2 | 0.222 | 0.250 |
|  | **CCG** | **38** | **45** | **3** | **8** | **0.079** | **0.178** |
| T | ACU | 17 | 22 | 2 | 0 | 0.118 | 0.000 |
|  | ACC | 34 | 47 | 8 | 12 | 0.235 | 0.255 |
|  | ACA | 3 | 1 | 2 | 0 | 0.667 | 0.000 |
|  | ACG | 6 | 7 | 1 | 3 | 0.167 | 0.429 |
| A | **GCU** | **19** | **28** | **1** | **7** | **0.053** | **0.250** |
|  | GCC | 9 | 11 | 2 | 3 | 0.222 | 0.273 |
|  | GCA | 22 | 33 | 5 | 5 | 0.227 | 0.152 |
|  | GCG | 28 | 52 | 4 | 6 | 0.143 | 0.115 |
| G | GGU | 68 | 85 | 6 | 6 | 0.088 | 0.071 |
|  | **GGC** | **35** | **29** | **4** | **9** | **0.114** | **0.310** |
|  | GGA | 0 | 0 | 0 | 0 | 0.000 | 0.000 |
|  | GGG | 3 | 1 | 1 | 0 | 0.333 | 0.000 |
| L | CUU | 6 | 3 | 0 | 2 | 0.000 | 0.667 |
|  | CUC | 15 | 7 | 3 | 0 | 0.200 | 0.000 |
|  | CUA | 0 | 0 | 0 | 0 | 0.000 | 0.000 |
|  | **CUG** | **100** | **125** | **6** | **13** | **0.060** | **0.104** |
| S | UCU | 23 | 24 | 2 | 1 | 0.087 | 0.042 |
|  | **UCC** | **31** | **27** | **9** | **2** | **0.290** | **0.074** |
|  | UCA | 0 | 1 | 0 | 0 | 0.000 | 0.000 |
|  | UCG | 3 | 5 | 0 | 0 | 0.000 | 0.000 |
| R | CGU | 61 | 75 | 6 | 5 | 0.098 | 0.067 |
|  | **CGC** | **28** | **24** | **10** | **1** | **0.357** | **0.042** |
|  | CGA | 1 | 0 | 0 | 0 | 0.000 | 0.000 |
|  | CGG | 0 | 0 | 0 | 0 | 0.000 | 0.000 |
| F | UUU | 11 | 9 | 1 | 1 | 0.091 | 0.111 |
|  | UUC | 33 | 26 | 2 | 2 | 0.061 | 0.077 |
| Y | UAU | 14 | 7 | 1 | 0 | 0.071 | 0.000 |
|  | UAC | 29 | 27 | 5 | 4 | 0.172 | 0.148 |
| H | CAU | 1 | 4 | 0 | 0 | 0.000 | 0.000 |
|  | **CAC** | **18** | **17** | **5** | **0** | **0.278** | **0.000** |
| Q | CAA | 8 | 2 | 1 | 0 | 0.125 | 0.000 |
|  | CAG | 50 | 48 | 3 | 2 | 0.060 | 0.042 |
| N | AAU | 3 | 1 | 0 | 1 | 0.000 | 1.000 |
|  | AAC | 48 | 47 | 4 | 4 | 0.083 | 0.085 |
| K | AAA | 56 | 62 | 3 | 3 | 0.054 | 0.048 |
|  | AAG | 24 | 25 | 2 | 3 | 0.083 | 0.120 |
| D | GAU | 30 | 34 | 2 | 2 | 0.067 | 0.059 |
|  | GAC | 62 | 47 | 8 | 7 | 0.129 | 0.149 |
| E | GAA | 89 | 83 | 2 | 3 | 0.022 | 0.036 |
|  | GAG | 33 | 26 | 1 | 3 | 0.030 | 0.115 |
| C | UGU | 5 | 7 | 0 | 0 | 0.000 | 0.000 |
|  | UGC | 2 | 8 | 1 | 3 | 0.500 | 0.375 |

**Supplementary table 4. Correlation study between TFD/FFD & synonymous ti/tv of whole gene/FFD**

|  |  |  |  | ti/tv (synonymous) | |  | Pearson R value |
| --- | --- | --- | --- | --- | --- | --- | --- |
| Genes | TFD | FFD | TFD/FFD | whole gene | FFD | whole gene/FFD |  |
| rpoB | 516 | 410 | 1.259 | 4.31 | 1.192 | 3.616 | **0.668** |
| rpoC | 480 | 482 | 0.996 | 2.929 | 1.441 | 2.032 |  |
| lacZ | 369 | 331 | 1.115 | 3.615 | 1.939 | 1.865 |  |
| lacY | 138 | 131 | 1.053 | 3.083 | 2.428 | 1.27 |  |
| kdpA | 133 | 223 | 0.596 | 3.686 | 3.091 | 1.192 |  |
| kdpB | 180 | 278 | 0.647 | 3.854 | 2.867 | 1.344 |  |
| araB | 182 | 219 | 0.831 | 4.917 | 3.733 | 1.317 |  |
| araA | 190 | 169 | 1.124 | 3.171 | 1.724 | 1.84 |  |
| bcsA | 279 | 275 | 1.015 | 4.344 | 3.4 | 1.278 |  |
| bcsB | 253 | 272 | 0.930 | 3.361 | 2.04 | 1.648 |  |

The correlation study between TFD:FFD ratio of all genes and synonymous ti/tv of whole gene/FFD was taken we got a strong positive Pearson r value as 0.668.

**Supplementary table 5. Ti/Tv ratio comparison of individual amino acids of four-fold degenerate codons between co-transcribed gene pairs.**

| Ti/Tv | rpoB | rpoC | lacZ | lacY | kdpA | kdpB | araB | araA | bcsA | bcsB |
| --- | --- | --- | --- | --- | --- | --- | --- | --- | --- | --- |
| Val | 1.00 | 0.60 | 1.57 | 1.50 | 6.00 | 2.29 | 2.75 | 1.43 | 2.50 | 1.67 |
| Pro | 2.00 | 2.67 | 2.00 | 2.00 | 2.40 | NA | 7.00 | 0.40 | 3.00 | 1.86 |
| Thr | 1.67 | 1.38 | 2.80 | 2.00 | 3.00 | 2.00 | 4.00 | 4.25 | 2.75 | 1.75 |
| Ala | 0.56 | 1.00 | 1.27 | 4.00 | 2.00 | 2.17 | 6.00 | 1.10 | 13.00 | 1.00 |
| Gly | 1.75 | 6.50 | 3.25 | 3.00 | 4.75 | 4.17 | 2.40 | 3.33 | 2.75 | NA |

Ti/Tv ratio comparison of four-fold degenerate amino acids individually in co-transcribed genes shows the difference in ratio even at amino acid level like the A in araB & araA is showing the difference, hence many such differences in ratio can be found at individual amino acid level. Genes showing NA values had zero transversions for the respective amino acids.

**Supplementary table 6. Codon count and mutation count of remaining 4 pairs of genes.**

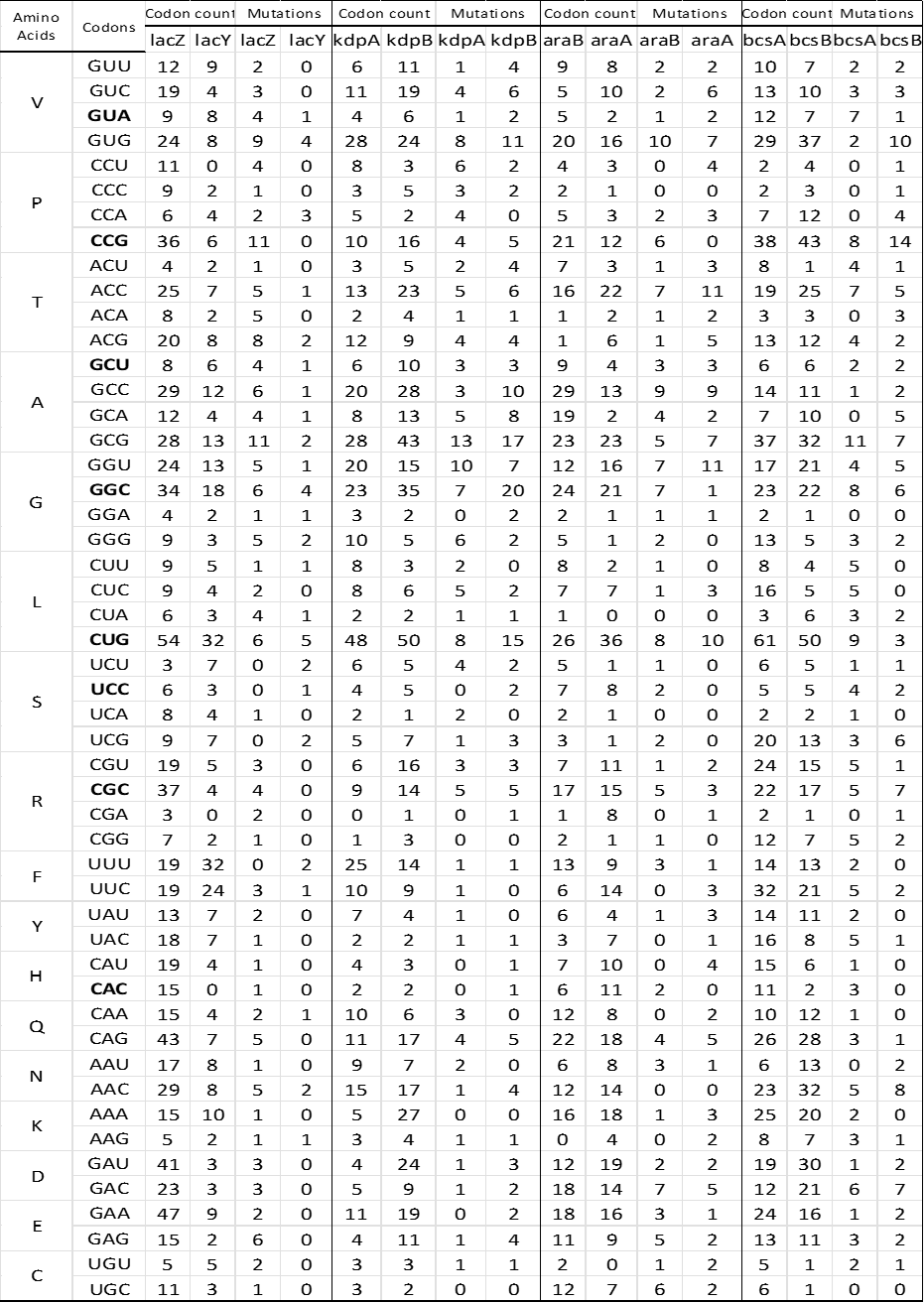

**Supplementary table 7. Strain wise mutation difference in co-transcribed genes**

Strainwise mutation difference in rpoB & rpoC

| Strains | Difference |
| --- | --- |
| 1 | -16 |
| 2 | -16 |
| 3 | -16 |
| 4 | -16 |
| 5 | -15 |
| 6 | -15 |
| 7 | -15 |
| 8 | -15 |
| 9 | -15 |
| 10 | -15 |
| 11 | -15 |
| 12 | -13 |
| 13 | -12 |
| 14 | -12 |
| 15 | -12 |
| 16 | -12 |
| 17 | -12 |
| 18 | -12 |
| 19 | -11 |
| 20 | -11 |
| 21 | -11 |
| 22 | -11 |
| 23 | -11 |
| 24 | -11 |
| 25 | -10 |
| 26 | -6 |
| 27 | -6 |
| 28 | -6 |
| 29 | -5 |
| 30 | -3 |
| 31 | -3 |
| 32 | -3 |
| 33 | -3 |
| 34 | -3 |
| 35 | -3 |
| 36 | -3 |
| 37 | -3 |
| 38 | -3 |
| 39 | -3 |
| 40 | -3 |
| 41 | -2 |
| 42 | -1 |
| 43 | -1 |
| 44 | 0 |
| 45 | 0 |
| 46 | 0 |
| 47 | 0 |
| 48 | 0 |
| 49 | 1 |
| 50 | 1 |
| 51 | 2 |
| 52 | 2 |
| 53 | 2 |
| 54 | 2 |
| 55 | 2 |
| 56 | 2 |
| 57 | 2 |
| 58 | 2 |
| 59 | 2 |
| 60 | 2 |
| 61 | 2 |
| 62 | 2 |
| 63 | 2 |
| 64 | 2 |
| 65 | 2 |
| 66 | 2 |
| 67 | 2 |
| 68 | 2 |
| 69 | 2 |
| 70 | 2 |
| 71 | 2 |
| 72 | 2 |
| 73 | 2 |
| 74 | 2 |
| 75 | 2 |
| 76 | 2 |
| 77 | 2 |
| 78 | 2 |
| 79 | 2 |
| 80 | 2 |
| 81 | 2 |
| 82 | 2 |
| 83 | 2 |
| 84 | 2 |
| 85 | 2 |
| 86 | 2 |
| 87 | 2 |
| 88 | 2 |
| 89 | 2 |
| 90 | 2 |
| 91 | 2 |
| 92 | 3 |
| 93 | 3 |
| 94 | 3 |
| 95 | 3 |
| 96 | 3 |
| 97 | 3 |
| 98 | 3 |
| 99 | 3 |
| 100 | 4 |
| 101 | 5 |
| 102 | 5 |
| 103 | 5 |
| 104 | 5 |
| 105 | 5 |
| 106 | 5 |
| 107 | 5 |
| 108 | 5 |
| 109 | 5 |
| 110 | 5 |
| 111 | 5 |
| 112 | 6 |
| 113 | 6 |
| 114 | 6 |
| 115 | 6 |
| 116 | 8 |
| 117 | 8 |
| 118 | 8 |
| 119 | 9 |
| 120 | 9 |
| 121 | 9 |
| 122 | 9 |
| 123 | 9 |
| 124 | 9 |
| 125 | 10 |
| 126 | 11 |
| 127 | 11 |
| 128 | 11 |
| 129 | 11 |
| 130 | 11 |
| 131 | 11 |
| 132 | 11 |
| 133 | 12 |
| 134 | 12 |
| 135 | 12 |
| 136 | 12 |
| 137 | 12 |
| 138 | 12 |
| 139 | 12 |
| 140 | 12 |
| 141 | 12 |
| 142 | 13 |
| 143 | 14 |
| 144 | 14 |
| 145 | 16 |
| 146 | 17 |
| 147 | 17 |
| 148 | 17 |
| 149 | 17 |
| 150 | 19 |

The individual strainwise mutation is shown in *rpoB* and *rpoC*. it indicates more number of indiivdual mutations are present in rpoC. Only 5 strains in both the genes share similar number of mutations out of 150 selected strains.

Strainwise mutation difference in kdpA & kdpB

|  | MUTATION DIFFERENCE BETWEEN TWO STRAINS |
| --- | --- |
| >NZ_CP010344 | -23 |
| >NZ_CP007265 | -22 |
| >NZ_CP007390 | -22 |
| >NZ_CP007391 | -22 |
| >NC_009800 | -21 |
| >NC_010468 | -19 |
| >NC_011415 | -19 |
| >NZ_CP006636 | -19 |
| >CP005998 | -19 |
| >NZ_CP009104 | -19 |
| >NZ_CP011018 | -19 |
| >NC_016902 | -18 |
| >NC_017635 | -18 |
| >NC_017660 | -18 |
| >NC_017664 | -18 |
| >NC_022364 | -18 |
| >NC_017641 | -15 |
| >NC_017663 | -15 |
| >NC_012892 | -14 |
| >NC_012967 | -14 |
| >NC_012971 | -14 |
| >NC_012947 | -14 |
| >NZ_CP010371 | -14 |
| >NZ_CP011938 | -14 |
| >NZ_CP014268 | -14 |
| >NZ_CP014269 | -14 |
| >NZ_AGTD01000001 | -13 |
| >NZ_CP009166 | -13 |
| >NZ_CP011134 | -13 |
| >NZ_CP013025 | -13 |
| >MG1655 | -12 |
| >NC_007779 | -12 |
| >NC_010473 | -12 |
| >NC_017625 | -12 |
| >NC_017633 | -12 |
| >NZ_AKBV01000001 | -12 |
| >NC_017638 | -12 |
| >NZ_AKVX01000001 | -12 |
| >NC_020518 | -12 |
| >CP006698 | -12 |
| >NZ_HG738867 | -12 |
| >NZ_CP008801 | -12 |
| >NZ_CP009273 | -12 |
| >NZ_CP009644 | -12 |
| >NZ_CP009789 | -12 |
| >NZ_CP009685 | -12 |
| >NZ_LM995446 | -12 |
| >NZ_LM993812 | -12 |
| >NZ_CP010438 | -12 |
| >NZ_CP010439 | -12 |
| >NZ_CP010440 | -12 |
| >NZ_CP010441 | -12 |
| >NZ_CP010442 | -12 |
| >NZ_CP010443 | -12 |
| >NZ_CP010444 | -12 |
| >NZ_CP010445 | -12 |
| >NZ_LN832404 | -12 |
| >NZ_CP007594 | -12 |
| >NZ_CP011320 | -12 |
| >NZ_CP011321 | -12 |
| >CP011322 | -12 |
| >CP011323 | -12 |
| >NZ_CP011324 | -12 |
| >NZ_CP011342 | -12 |
| >NZ_CP011343 | -12 |
| >NZ_CP011495 | -12 |
| >NZ_CP012125 | -12 |
| >NZ_CP012126 | -12 |
| >NZ_CP012127 | -12 |
| >NZ_CP011113 | -12 |
| >NZ_CP012868 | -12 |
| >NZ_CP012869 | -12 |
| >NZ_CP012870 | -12 |
| >NZ_CP013253 | -12 |
| >NZ_CP014225 | -12 |
| >NZ_CP014270 | -12 |
| >NZ_CP014272 | -12 |
| >NC_020163 | -12 |
| >NZ_CP010315 | -12 |
| >NZ_CP007442 | -12 |
| >NZ_CP013112 | -12 |
| >NZ_CP007491 | -12 |
| >NC_009801 | -10 |
| >NZ_CP007592 | -10 |
| >NZ_CP009106 | -9 |
| >NC_011748 | -8 |
| >NC_011741 | -8 |
| >NC_018650 | -8 |
| >NC_018658 | -8 |
| >NC_018661 | -8 |
| >NZ_HF572917 | -8 |
| >NZ_CP011331 | -8 |
| >NZ_CP011416 | -7 |
| >NC_013364 | -6 |
| >NC_013361 | -6 |
| >NZ_CP006262 | -2 |
| >NZ_CP007133 | -2 |
| >NZ_CP010585 | -1 |
| >NZ_CP007392 | -1 |
| >NZ_CP007393 | -1 |
| >NZ_CP007394 | -1 |
| >NZ_CP006027 | -1 |
| >NZ_CP007136 | -1 |
| >NC_010498 | 2 |
| >NC_013941 | 3 |
| >NC_017656 | 3 |
| >NZ_CP008805 | 3 |
| >NC_002695 | 4 |
| >NC_011353 | 4 |
| >NC_013008 | 4 |
| >NC_017906 | 4 |
| >NZ_CP008957 | 4 |
| >NZ_CP010304 | 4 |
| >NZ_CP012802 | 4 |
| >NZ_CP014314 | 4 |
| >NC_013654 | 4 |
| >NC_017644 | 4 |
| >NZ_HG941718 | 4 |
| >NZ_CP010876 | 4 |
| >NZ_CP013658 | 4 |
| >NZ_CP013831 | 4 |
| >NZ_CP013835 | 4 |
| >NZ_CP006632 | 5 |
| >NC_011993 | 8 |
| >NC_004431 | 9 |
| >NC_017651 | 9 |
| >NC_017652 | 9 |
| >NZ_CP007799 | 9 |
| >NZ_CP009072 | 9 |
| >NC_017631 | 10 |
| >NZ_CP014197 | 12 |
| >NZ_CP013029 | 12 |
| >NC_011750 | 16 |
| >NC_017646 | 16 |
| >NC_008253 | 21 |
| >NZ_CP005930 | 24 |
| >NC_007946 | 25 |
| >NC_008563 | 25 |
| >NC_017628 | 25 |
| >CU928161 | 25 |
| >NC_017632 | 25 |
| >NC_022370 | 25 |
| >NZ_CP007149 | 25 |
| >NZ_CP012635 | 25 |
| >NZ_CP012625 | 25 |
| >NZ_CP012633 | 25 |
| >NZ_CP012631 | 25 |
| >NZ_CP009859 | 26 |
| >NZ_CP008697 | 26 |
| >NC_017626 | 29 |
| >NZ_CP010816 | 30 |

The individual strainwise mutation is shown in *kdpA* and *kdpB*. it indicates more number of indiivdual mutations are present in *kdpA*. There were no such strains having similar mutation sin both the co-transcribed genes. hence it shows the difference in mutations in both the co-transcribed genes is present even in intra species comparison.
